# Genozip 14 - advances in compression of BAM and CRAM files

**DOI:** 10.1101/2022.09.12.507582

**Authors:** Divon Lan, Bastien Llamas

**Author notes:** Corresponding author: DL.

## Abstract

**Summary:** Genozip performs compression of a wide range of genomic data, including widely used FASTQ, BAM and VCF file formats. Here, we introduce the latest advancement in Genozip technology, focused on compression of BAM and CRAM files. We demonstrate Genozip’s ability to compress data generated by a variety of study types (e.g., whole genome sequencing, DNA methylation, RNASeq), sequencing technologies and aligners, up to 2.7 times better than the current state of the art compressor, CRAM version 3.1.

**Availability and implementation:** Genozip is freely available for academic research use and has been tested for Linux, Mac and Windows. Installation instructions are available at https://genozip.com/installing.html. A user manual is available at https://genozip.com/manual.html.

**Supplementary information:** Supplementary data are available.

## 1 Introduction

The rapid accumulation of genomic data is a growing problem, driven by the compounded effects of a continuing drop in sequencing costs and an increase in clinical and research applications of genomics. It is not surprising then that the field of genomic data compression is an area of active research, focused mostly on the three file formats that constitute the bulk of the accumulated genomic data - namely, FASTQ (e.g., (Chandak *et al*., 2019), (Dufort Y Álvarez *et al*., 2020)), SAM (e.g., (Bonfield, 2022), (Hach *et al*., 2014)) and VCF (e.g., (Lan *et al*., 2020), (Deorowicz *et al*., 2021)). For SAM data, the compressed format most widely used is BAM, which implements compression based on the gzip standard (Li *et al*., 2009), while CRAM is an emerging format based on significantly better, modern codecs. Genozip (Lan *et al*., 2021) is a software system for compression of all the abovementioned and several other types of genomic files (FASTA, GFF3, GVF, 23andMe and others). Here, we introduce Genozip version 14, which focuses on the advancement of compression of SAM / BAM / CRAM files. We show that Genozip 14 compresses significantly better than both the current default version of CRAM, 3.0, as well as its recently introduced newer version, 3.1.

## 2 Software description

Genozip is an easy to use command-line tool, which consists of four commands: `genozip, genounzip, genols` and `genocat`. The latter is a tool for accessing and subsetting data compressed with Genozip, allowing efficient use of Genozip-compressed files in bioinformatics pipelines. Genozip is a feature-rich tool with many capabilities not covered in this article focused on BAM/CRAM compression - see the user manual for more information.

The key difference between the Genozip and CRAM approaches lies in the product philosophy itself. Since CRAM aims to be an official standard (CRAM), its development process is driven by a slow, consensus-oriented, multi-organisation collaboration, and it is purposely oblivious to the many non-standard extensions of SAM tags introduced by tools developed to support various study types, including DNA methylation, RNAseq, single-cell analysis, and metagenomics. Genozip, in contrast, focuses on delivering the best compression for any emerging laboratory methods, sequencing technologies, novel SAM tags and study types, making it much more flexible and versatile than other compressors. Genozip employs specific methods for compressing 86 standard and non-standard tags generated by 28 commonly used aligners and other SAM/BAM-producing software packages (Table S1), as well as generic methods for compressing all other tags. In addition, Genozip has specific methods to compress data generated by sequencing technologies such as Illumina, Pacific Biosciences, Oxford Nanopore, MGI and Ion Xpress.

## 3 Benchmark

We benchmarked Genozip 14 against two versions of CRAM: CRAM 3.0, which is the default version in samtools, and CRAM 3.1, the latest CRAM version, for which the current version of samtools (1.15.1) produces the warning “*This is a technology demonstration that should not be used for archival data*”. We acknowledge that other BAM compression systems exist, such as DeeZ (Hach *et al*., 2014) and MPEG-G (Voges *et al*., 2021). However, they have been shown to be significantly inferior to both CRAM 3.1 and Genozip 13 (Bonfield, 2022), therefore we did not include them in our benchmark.

For the sake of reproducibility and suitable comparison, we used the same three benchmark files (2 BAM files and one CRAM file) from (Bonfield, 2022). We also included 11 additional BAM files that represent a wide variety of aligners, sequencing technologies and study types. More details about the benchmark process, including file sources (Table S2) and reference files (Table S3), can be found in the Supplementary Information.

We tested each of the 14 files using each of the 3 tools (CRAM 3.0, CRAM 3.1 and Genozip 14) in the default (“*normal*”) mode, as well as in the highest available lossless compression (“*best*”) mode, thereby producing 28 unique benchmarks (Table S4; Figures 1 and S1–S28). When comparing Genozip 14 to CRAM 3.1 in their respective *normal* and *best* modes, our results show that Genozip compressed better than CRAM in 27 of these 28 benchmarks (Table S4; Figures 1 and S1–S10, S12–S14). Genozip in *best* mode performed marginally worse than CRAM 3.1 in *best* mode for the total RNA-Seq dataset T11 (Table S4; Figure S11). The compression ratio advantage of Genozip 14 varied widely: 2.1–13.1X better when compared to BAM and 1.1–2.5X better when compared to CRAM 3.1 in *normal* mode, and 2.1X–15.9X vs BAM and 1X–2.7X vs CRAM 3.1 in *best* mode (Table S4; Figure 1). We observed that Genozip’s advantage over CRAM grows with the complexity of the data. For example, files with secondary or supplementary alignments (e.g., T4), files with bisulfite-treated reads (e.g., T14), files with barcodes (e.g., T12), and files produced from long reads (e.g., T6, 7) benefited substantially from Genozip compression.

**Figure 1:**
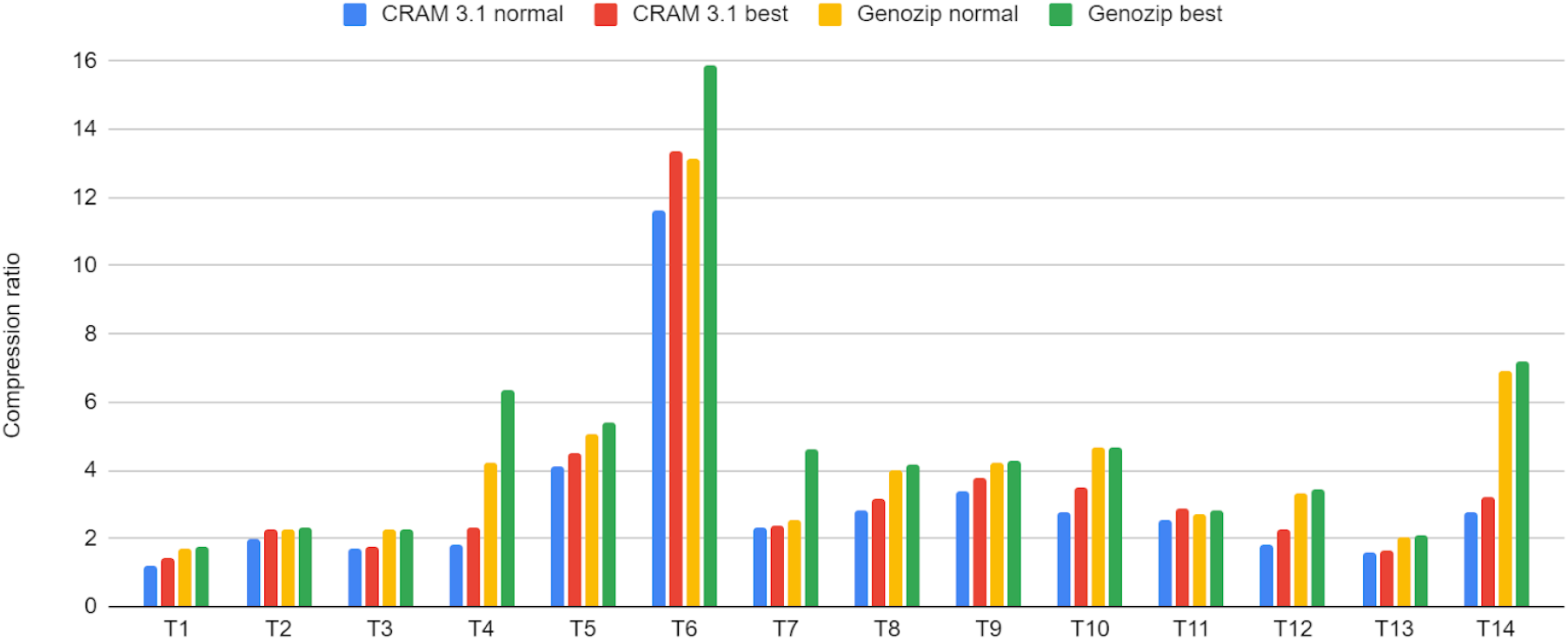
benchmark of compression ratio using 14 files (T1 through T14). T1 is a CRAM 3.0 file while T2 to T14 are BAM files. The Y axis is the compression ratio achieved compared to the original CRAM 3.0 (T1) or BAM (T2-14) files. We performed four tests for each file: CRAM 3.1 in normal mode, CRAM 3.1 in best mode, Genozip 14 in normal mode and Genozip 14 in best mode. The benchmark files represent a wide range of study types: **T1**: WGS - NovaSeq file used in (Bonfield, 2022); **T2**: WGS - HiSeq 2000 file used in (Bonfield, 2022); **T3**: WGS - PacBio CLR file used in (Bonfield, 2022); **T4**: RNA-seq - transcriptome alignments; **T5**: RNA-seq - genome alignments; **T6**: Long read RNA-seq; **T7**: WGS - GIAB; **T8**: DNase-Seq; **T9**: STARR-seq; **T10**: scRNA-seq; **T11**: totalRNA-seq; **T12**: 1:1 Mixture of Fresh Frozen Human and Mouse Cells; **T13**: WGS - Nanopore MinION; **T14**: WGBS paired-end (methylation). See Supplementary Information for more details on the test files, compression and decompression times, and comparison to CRAM 3.0.

Genozip 14 compressed faster than CRAM in 19 of the 28 cases (Table S5). Genozip 14 decompressed faster than CRAM in 4 cases, slower in 22 cases, whereas in the 2 remaining cases CRAM failed to decompress the file (Table S5). Moreover, when comparing CRAM in *best* mode to Genozip in *normal* mode, Genozip still compressed better than CRAM in 12 of the 14 tests (Table S4; Figures 1 and S1–S5, S7–S10, S12–S14), and faster than CRAM in all tests (Table S5).

## 4 Conclusion

Genozip 14 demonstrates significantly superior compression of BAM and CRAM files compared to CRAM 3.1, and hence it would be a good choice for users seeking to minimise consumption of storage resources, for both archival purposes and for use in bioinformatics pipelines.

## Supporting information

Genozip 14 - Supplementary Information

## Acknowledgements

We thank Felix Krueger for his support with Bismark, Weilong Guo for this support with

BS-Seeker2, Colin Farrell and Wenbin Guo for their support with BSBolt, 10xGenomics for their support with cellranger and longranger, Qiagen for their support with CLC Genomics Workbench, Colin Hercus for his support with Novoalign, BGI for their support with MGI data, and James Bonfield for his support with CRAM as well as his kindness and encouragement.

