## Supplementary material for "Genozip 14 - advances in compression of BAM and CRAM files": Genozip 14 - Supplementary Information

Divon Lan[1,*], Bastien Llamas[1,2,3,4]

[1] Australian Centre for Ancient DNA, School of Biological Sciences, The Environment Institute, Faculty of Sciences, The University of Adelaide, Adelaide SA 5005, Australia
[2] Centre of Excellence for Australian Biodiversity and Heritage (CABAH), School of Biological Sciences, University of Adelaide, Adelaide, SA 5005, Australia
[3] National Centre for Indigenous Genomics, Australian National University, Canberra, ACT 0200, Australia
[4] Telethon Kids Institute, Adelaide, SA 5000, Australia

**Custom methods and codecs** **3**

**Benchmark setup** **4**

Benchmark files 4

Benchmark implementation details 7

**Benchmark results** **8**

### Custom methods and codecs

Genozip 14 contains methods and codecs for compressing non-standard fields or data with specific characterisation generated by third party aligners and other software packages.

Table S1: SAM/BAM-producing software packages, which generate non-standard SAM tags, for which Genozip 14 contains specific codecs or compression methods

| | |
|---|---|
| BWA | TMap |
| minimap2 | HISAT2 |
| bowtie | NovoCraft - NovoAlign and NovoSort |
| bowtie2 | Razer3 |
| STAR | Blasr |
| Illumina Dragen | NGMLR |
| Gem3 | Delve |
| gem-2-sam | TopHat |
| Bismark | CPU |
| BS-Seeker2 | 10xGenomics CellRanger |
| Winnowmap | 10xGenomics LongRanger |
| PacBio baz2bam | CLC Genomics Workbench |
| BBMap | BSBolt |
| GATK - BQSR and MergeBamAlignment | biobambam - bamsort and bamclipXT |

### Benchmark setup

#### Benchmark files

Table S2: List of benchmark files

| ID | Filename | Study type | Sequencer | Aligner | Data URL | Citation |
|---|---|---|---|---|---|---|
| T1 | NA12878.final.cram | WGS - NovaSeq file used in (Bonfield, 2022) | Illumina NovaSeq | bwa | ftp://ftp.sra.ebi.ac.uk/vol1/run/ERR323/ERR3239334/NA12878.final.cram | (The 1000 Genomes Project Consortium, 2012) |
| T2 | NA12878_S1.bam | WGS - HiSeq 2000 file used in (Bonfield, 2022) | Illumina HiSeq 2000 | bwa | ftp://ftp.sra.ebi.ac.uk/vol1/run/ERR194/ERR194147/NA12878_S1.bam | (The 1000 Genomes Project Consortium, 2012) |
| T3 | NA12878.pacbio.bwa-sw.20140202.bam | WGS - PacBio CLR - file used in (Bonfield, 2022) | PacBio | bwa sw | ftp://ftp.1000genomes.ebi.ac.uk/vol1/ftp/technical/working/20131209_na12878_pacbio/si/NA12878.pacbio.bwa-sw.20140202.bam | (The 1000 Genomes Project Consortium, 2012) |
| T4 | ENCFF047UEJ.bam | RNA-seq - transcriptome alignments | Illumina HiSeq 2500 | STAR | https://www.encodeproject.org/files/ENCFF047UEJ/@@download/ENCFF047UEJ.bam | (de Souza, 2012) |
| T5 | ENCFF575KZB.bam | RNA-seq - genome alignments | Illumina HiSeq 2500 | STAR | https://www.encodeproject.org/files/ENCFF575KZB/@@download/ENCFF575KZB.bam | (de Souza, 2012) |
| T6 | ENCFF900XHI.bam | long read RNA-seq | PacBio Sequel II | minimap2 | https://www.encodeproject.org/files/ENCFF900XHI/@@download/ENCFF900XHI.bam | (de Souza, 2012) |
| T7 | sorted_final_merged.bam | WGS (GIAB) | PacBio | blasr | ftp://ftp-trace.ncbi.nlm.nih.gov:21/giab/ftp/data/NA12878/NA12878_PacBio_MtSinai/sorted_final_merged.ba | (Zook, 2012) |

| | | | | | | |
|---|---|---|---|---|---|---|
| | | | | | m | |
| T8 | ENCFF069XDC.bam | DNase-Seq | Illumina HiSeq 4000 | bwa sampe | https://www.encodeproject.org/files/ENCFF069XDC/@@download/ENCFF069XDC.bam | (de Souza, 2012) |
| T9 | ENCFF669HBS.bam | STARR-seq | Illumina NovaSeq 6000 | bowtie2 | https://www.encodeproject.org/files/ENCFF669HBS/@@download/ENCFF669HBS.bam | (de Souza, 2012) |
| T10 | ENCFF046VPK.bam | scRNA-seq | Illumina NextSeq 2000 | STARsolo | https://www.encodeproject.org/files/ENCFF046VPK/@@download/ENCFF046VPK.bam | (de Souza, 2012) |
| T11 | ENCFF460RWK.bam | totalRNA-seq | Illumina HiSeq 2500 | STAR | https://www.encodeproject.org/files/ENCFF460RWK/@@download/ENCFF460RWK.bam | (de Souza, 2012) |
| T12 | hgmm_10k_v3_possorted_genome_bam.bam | Single cell - 1:1 Mixture of Fresh Frozen Human and Mouse Cells | Illumina NovaSeq | STAR + cellranger | https://www.10xgenomics.com/resources/datasets/10-k-1-1-mixture-of-fresh-frozen-human-hek-293-t-and-mouse-nih-3-t-3-cells-v-3-chemistry-3-standard-3-0-0 | N/A |
| T13 | ENCFF283TLK.bam | WGS - Nanopore MinION | Oxford Nanopore MinION | ngmlr | https://www.encodeproject.org/files/ENCFF283TLK/@@download/ENCFF283TLK.bam | (de Souza, 2012) |
| T14 | ENCFF786GJA.bam | WGBS paired-end (Methylation) | Illumina HiSeq X Ten | Bismark | https://www.encodeproject.org/files/ENCFF786GJA/@@download/ENCFF786GJA.bam | (de Souza, 2012) |

Table S3: Reference files used

| ID | Details | Used for | URL |
|---|---|---|---|
| grch38 | | T1,T9 | ftp://ftp.1000genomes.ebi.ac.uk/vol1/ftp/technical/reference/GRCh38_reference_genome/GRCh38_full_analysis_set_plus_decoy_hla.fa |
| gca | | T8,T13,T14 and generating gca_ercc | http://www.epigenomes.ca/data/CEMT/resources/GCA_000001405.15_GRCh38_no_alt_analysis_set.fna.gz/scratch4/mschatz1/mkirsche/GCA_000001405.15_GRCh38_no_alt_analysis_set.fna |
| ercc | | Generating gca_ercc, mm10_ercc | https://www-s.nist.gov/srmors/view_datafiles.cfm?srm=2374 |
| phiX | | Generating gca_ercc, mm10_ercc | https://www.ncbi.nlm.nih.gov/nuccore/NC_001422.1?report=fasta |
| gca_ercc | Created by concatenating gca, ercc and phiX | T5,T6,T11 | |
| mm10 | | Generating mm10_ercc | https://www.encodeproject.org/files/mm10_no_alt_analysis_set_ENCODE/@@download/mm10_no_alt_analysis_set_ENCODE.fasta.gz |
| mm10_ercc | Created by concatenating mm10, ercc and phiX | T10 | |
| hg19 | | T2,T7 | http://hgdownload.cse.ucsc.edu/goldenpath/hg19/bigZips/hg19.fa.gz |
| hs37d5 | | T3 | ftp://ftp.1000genomes.ebi.ac.uk/vol1/ftp/technical/reference/phase2_reference_assembly_sequence |
| NONE | These files were compressed without a reference since the reference with which the BAM was generated was not readily available | T4,T12 | |

#### Benchmark implementation details

For Genozip, we used version 14.0.0.
*Normal* mode: the options used were `--no-test`, and `--reference` in cases where a reference was used (see table S3).
*Best* mode: the options used were `--best`, `--no-test`, and `--reference` in cases where a reference was used (see table S3).

For CRAM 3.0, we used samtools version 1.15.1.
*Normal* mode: the options used were `-@61 -O cram`, and `-T` in cases where a reference was used (see table S3).
*Best* mode: the options used were `-@61 -O cram,archive,level=9,use_lzma,use_bzip2`, and `-T` and `--reference` in cases where a reference was used (see table S3), and an additional `-O` sub-option `no_ref` if not.

For CRAM 3.1, we used the same as CRAM 3.0, but with an additional `-O` sub-option `version=3.1`.

The tests were conducted on a Linux machine with 56 cores.

Note: Genozip is able to compress CRAM files (for example, T1). When compressing, it invokes `samtools` to read the file as BAM, and when decompressing, it outputs the file in BAM format.

Note on reference files: preparation and indexing of the reference files for CRAM, and preparation and generation of the reference cache files in Genozip (which happens upon first use of a reference file in Genozip) were done ahead of running the benchmarks, and hence are not included in the time measurements.

Note on file T3: in (Bonfield, 2022) the author subsets the file to remove all secondary alignments and auxiliary tags AS:i, PG:Z, SA:Z, XS:i. This is done for ease of comparison with MPEG-G. In our benchmark, we compress the entire file.

Note on files T5, T6, T10, T11: These files contain ERCC contigs, and the original reference file used to create these files is not available. We did our best to recreate the reference file (see Table S3) however, 8 of the 96 ERCC contigs have lengths that mismatch the lengths specified in the SAM header. Nevertheless, both Genozip and CRAM are able to compress these files without issues despite these mismatches.

Note on file T10: CRAM hangs when trying to decompress the file. This happens in both versions 3.0 and 3.1 and in both *normal* and *best* modes.

Note of files T4 and T12: the original reference files used to generate these BAM files are not available and are not trivial to recreate. We therefore compare Genozip and CRAM in their respective reference-free modes in these cases.

Note on threads: Genozip over-subscribes threads to cores - setting the limit at 1.1 threads per core, or 61 threads on our test machine. samtools is limited by default to a single thread. For a fair comparison, we use the `-@61` command line option to allow samtools to use up to 61 threads as well.

Note: By default, Genozip tests every file after compression by decompressing it in memory and comparing the Adler32 signature of the original and reconstructed files. CRAM does not provide this functionality. For a fair comparison, we use `--no-test` to disable this.

### Benchmark results

Table S4: Benchmark results - compression ratio

| ID | BAM | CRAM 3.0 normal | | CRAM 3.0 best | | CRAM 3.1 normal | | CRAM 3.1 best | | Genozip normal | | | | Genozip best | | |
|---|---|---|---|---|---|---|---|---|---|---|---|---|---|---|---|---|
| | size | size | vs BAM | size | vs BAM | size | vs BAM | size | vs BAM | size | vs BAM | vs CRAM 3.1 normal | vs CRAM 3.1 best | size | vs BAM | vs CRAM 3.1 best |
| T1 | 15797182294 | 15057033571 | 1.05* | 12875681480 | 1.23* | 12866586165 | 1.23* | 11274600408 | 1.40* | 9334566499 | 1.69* | 1.38 | 1.21 | 9015105386 | 1.75* | 1.25 |
| T2 | 121691186161 | 65758948961 | 1.85 | 62846035154 | 1.94 | 61871207684 | 1.97 | 53816301380 | 2.26 | 53261807466 | 2.28 | 1.16 | 1.01 | 52005672886 | 2.34 | 1.03 |
| T3 | 57785894776 | 34142797882 | 1.69 | 33134668135 | 1.74 | 34012686431 | 1.70 | 32931470457 | 1.75 | 25777141400 | 2.24 | 1.32 | 1.28 | 25442443993 | 2.27 | 1.29 |
| T4 | 2065298931 | 1191378160 | 1.73 | 896039669 | 2.30 | 1150818566 | 1.79 | 888342282 | 2.32 | 485676014 | 4.25 | 2.37 | 1.83 | 324674984 | 6.36 | 2.74 |
| T5 | 2322825802 | 703852124 | 3.30 | 572843075 | 4.05 | 560935284 | 4.14 | 515482780 | 4.51 | 458320451 | 5.07 | 1.22 | 1.12 | 428436333 | 5.42 | 1.20 |
| T6 | 1308956918 | 117706032 | 11.12 | 96229390 | 13.60 | 112853011 | 11.60 | 97959482 | 13.36 | 99830250 | 13.11 | 1.13 | 0.98 | 82497181 | 15.87 | 1.19 |
| T7 | 146870854017 | 64230408277 | 2.29 | 62986763535 | 2.33 | 63471703737 | 2.31 | 62370839534 | 2.35 | 58277679349 | 2.52 | 1.09 | 1.07 | 31796498533 | 4.62 | 1.96 |
| T8 | 6668114400 | 2569641796 | 2.59 | 2244070944 | 2.97 | 2361434390 | 2.82 | 2107236567 | 3.16 | 1660198748 | 4.02 | 1.42 | 1.27 | 1598034373 | 4.17 | 1.32 |
| T9 | 5322159747 | 1732874059 | 3.07 | 1558317263 | 3.42 | 1560269168 | 3.41 | 1407535434 | 3.78 | 1258863510 | 4.23 | 1.24 | 1.12 | 1242102254 | 4.28 | 1.13 |
| T10 | 1306491967 | 517065206 | 2.53 | 395166766 | 3.31 | 472757390 | 2.76 | 375053549 | 3.48 | 280170704 | 4.66 | 1.69 | 1.34 | 277976393 | 4.70 | 1.35 |
| T11 | 4171989444 | 1865069285 | 2.24 | 1748221844 | 2.39 | 1638636424 | 2.55 | 1449031061 | 2.88 | 1544543630 | 2.70 | 1.06 | 0.94 | 1473370983 | 2.83 | 0.98 |
| T12 | 53694972688 | 32965299374 | 1.63 | 24752933445 | 2.17 | 29550304054 | 1.82 | 23951541634 | 2.24 | 16012039113 | 3.35 | 1.85 | 1.50 | 15552234159 | 3.45 | 1.54 |
| T13 | 222354403768 | 138092782957 | 1.61 | 137253032579 | 1.62 | 137969266061 | 1.61 | 136579234155 | 1.63 | 108704693272 | 2.05 | 1.27 | 1.26 | 107386919618 | 2.07 | 1.27 |
| T14 | 198026253992 | 72250800865 | 2.74 | 63240712720 | 3.13 | 70809231816 | 2.80 | 61730961113 | 3.21 | 28621048627 | 6.92 | 2.47 | 2.16 | 27448746040 | 7.21 | 2.25 |

* T1 is a CRAM 3.0 file, hence the ratio shown is vs this CRAM 3.0 file, not vs BAM.

#### Table S5: Benchmark results - execution time

| ID | CRAM 3.0 normal | | CRAM 3.0 best | | CRAM 3.1 normal | | CRAM 3.1 best | | Genozip normal | | Genozip best | |
|---|---|---|---|---|---|---|---|---|---|---|---|---|
| | **compression** | **decompression** | **compression** | **decompression** | **compression** | **decompression** | **compression** | **decompression** | **compression** | **decompression** | **compression** | **decompression** |
| T1 | real 10m44.849s<br>user 70m16.331s<br>sys 2m42.902s | real 4m46.950s<br>user 68m14.161s<br>sys 3m7.279s | real 40m16s | real 6m:08s | real 9m28.712s<br>user 70m39.559s<br>sys 2m34.316s | real 4m59.196s<br>user 67m19.242s<br>sys 3m15.959s | real 31m2.550s<br>user 924m15.612s<br>sys 130m14.398s | real 5m56.571s<br>user 107m5.768s<br>sys 6m1.357s | real 13m29.423s<br>user 161m34.085s<br>sys 25m7.138s | real 15m26.880s<br>user 141m34.885s<br>sys 28m31.453s | real 14m31.356s<br>user 237m7.512s<br>sys 30m10.577s | real 15m32.131s<br>user 159m15.951s<br>sys 18m18.474s |
| T2 | elapsed 13:10.27<br>user 6013.17<br>system 306.05 | real 8m29.337s<br>user 142m12.332s<br>sys 5m45.558s | elapsed 45:27.51<br>user 117554.65<br>system 8064.20 | real 23m23.922s<br>user<br>157m46.992s<br>sys 8m24.320s | real 12m32.251s<br>user<br>125m39.067s<br>sys 7m15.862s | real 8m53.323s<br>user 156m22.005s<br>sys 6m26.969s | real 29m31.013s<br>user 999m44.409s<br>sys 88m59.194s | real 10m59.384s<br>user 276m4.841s<br>sys 8m59.176s | real 28m35.322s<br>user 1113m30.524s<br>sys 61m28.963s | real 24m22.641s<br>user 727m52.240s<br>sys 34m19.957s | real 32m31.366s<br>user 1372m17.330s<br>sys 63m4.126s | real 22m39.016s<br>user 942m41.720s<br>sys 42m11.937s |
| T3 | real 2m21.520s<br>user 28m21.364s<br>sys 3m13.433s | elapsed 2:13.77<br>user 3521.40<br>system 138.67 | elapsed 30:55.89<br>user 88148.25<br>system 2758.61 | elapsed 3:32.70<br>user 3663.14<br>system 297.46 | real 2m17.767s<br>user 31m11.869s<br>sys 2m40.770s | real 2m37.711s<br>user 57m53.222s<br>sys 2m44.889s | real 38m38.064s<br>user 1822m40.106s<br>sys 54m37.537s | real 4m15.433s<br>user 108m2.575s<br>sys 5m57.736s | real 6m12.762s<br>user 148m54.013s<br>sys 12m32.025s | real 8m59.801s<br>user 194m57.279s<br>sys 12m50.004s | real 9m18.720s<br>user 210m3.226s<br>sys 15m21.433s | real 10m54.153s<br>user 222m18.630s<br>sys 14m54.031s |
| T4 | real 0m18.349s<br>user 4m55.773s<br>sys 0m8.816s | real 1m5.516s<br>user 7m0.112s<br>sys 45m38.904s | real 4m11.431s<br>user 200m45.894s<br>sys 6m42.566s | real 1m7.942s<br>user 8m2.963s<br>sys 45m24.731s | real 0m18.516s<br>user 5m36.996s<br>sys 0m10.639s | real 1m52.146s<br>user 7m40.453s<br>sys 52m46.329s | real 4m52.396s<br>user 233m38.068s<br>sys 7m59.858s | real 1m13.626s<br>user 9m0.944s<br>sys 48m24.314s | real 1m52.474s<br>user 15m37.209s<br>sys 3m37.978s | real 1m51.080s<br>user 6m3.839s<br>sys 1m26.179s | real 1m31.169s<br>user 14m28.722s<br>sys 3m29.721s | real 1m5.596s<br>user 7m14.021s<br>sys 0m54.872s |
| T5 | real 0m48.909s<br>user 3m29.952s<br>sys 0m24.392s | real 1m33.537s<br>user 3m35.018s<br>sys 0m23.529s | real 3m8.049s<br>user 95m19.033s<br>sys 6m36.426s | real 0m31.872s<br>user 4m51.371s<br>sys 0m38.159s | real 0m49.452s<br>user 4m11.557s<br>sys 0m36.910s | real 0m19.713s<br>user 3m27.300s<br>sys 0m24.560s | real 2m57.345s<br>user 106m31.005s<br>sys 7m51.119s | real 0m35.557s<br>user 4m36.219s<br>sys 1m16.041s | real 0m32.891s<br>user 13m53.089s<br>sys 2m16.651s | real 0m37.955s<br>user 6m45.043s<br>sys 0m38.627s | real 0m57.270s<br>user 11m5.932s<br>sys 1m47.792s | real 0m36.624s<br>user 7m26.463s<br>sys 0m34.945s |
| T6 | real 1m7.162s<br>user 12m5.502s<br>sys 3m27.353s | real 0m14.806s<br>user 1m43.974s<br>sys 0m53.367s | real 0m41.963s<br>user 1m19.903s<br>sys 0m13.189s | real 0m10.326s<br>user 1m18.695s<br>sys 0m19.797s | real 0m42.753s<br>user 1m29.412s<br>sys 0m12.496s | real 0m21.419s<br>user 1m11.847s<br>sys 0m33.400s | real 1m13.058s<br>user 16m26.650s<br>sys 2m43.949s | real 0m14.850s<br>user 1m38.301s<br>sys 0m58.673s | real 0m6.466s<br>user 2m35.437s<br>sys 0m16.071s | real 0m24.773s<br>user 4m26.430s<br>sys 0m24.635s | real 0m18.035s<br>user 2m28.421s<br>sys 0m12.615s | real 0m24.928s<br>user 3m17.380s<br>sys 0m12.813s |
| T7 | elapsed 7:57.24<br>user 5819.20<br>system 453.16 | elapsed 11:28.97<br>user 12536.38<br>system 560.71 | elapsed 2:02:01<br>user 317653.43<br>system 19214.82 | real 19m0.753s<br>user<br>241m55.294s<br>sys 26m57.759s | real 8m53.362s<br>user 86m6.034s<br>sys 11m24.832s | real 11m25.616s<br>user 199m25.359s<br>sys 9m52.199s | real 83m47.354s<br>user 3163m0.696s<br>sys 150m45.401s | real 17m51.432s<br>user<br>284m51.183s<br>sys 24m18.447s | real 23m41.183s<br>user 1004m26.418s<br>sys 59m9.421s | real 23m37.774s<br>user 439m47.490s<br>sys 37m59.990s | real 42m13.043s<br>user 755m11.202s<br>sys 69m37.955s | real 38m5.082s<br>user 704m21.097s<br>sys 54m52.166s |
| T8 | real 3m27.705s<br>user 20m4.741s<br>sys 16m39.244s | real 1m10.329s<br>user 15m5.220s<br>sys 28m14.046s | real 6m12.669s<br>user 174m50.396s<br>sys 24m35.008s | real 1m14.659s<br>user 14m50.047s<br>sys 20m2.487s | real 2m45.430s<br>user 17m9.024s<br>sys 20m33.755s | real 1m56.739s<br>user 17m39.608s<br>sys 53m51.478s | real 5m41.257s<br>user 179m6.975s<br>sys 26m16.166s | real 1m33.932s<br>user 20m24.501s<br>sys 41m29.958s | real 1m25.422s<br>user 48m8.419s<br>sys 3m2.816s | real 1m4.617s<br>user 20m9.893s<br>sys 1m20.277s | real 1m42.099s<br>user 63m0.803s<br>sys 2m30.331s | real 1m32.029s<br>user 33m53.611s<br>sys 1m48.843s |

| ID | CRAM 3.0 normal | | CRAM 3.0 best | | CRAM 3.1 normal | | CRAM 3.1 best | | Genozip normal | | Genozip best | |
|---|---|---|---|---|---|---|---|---|---|---|---|---|
| | **compression** | **decompression** | **compression** | **decompression** | **compression** | **decompression** | **compression** | **decompression** | **compression** | **decompression** | **compression** | **decompression** |
| T9 | real 0m55.191s<br>user 5m49.564s<br>sys 0m30.356s | real 0m30.894s<br>user 8m18.977s<br>sys 0m31.140s | real 5m45.064s<br>user 259m27.186s<br>sys 11m1.915s | real 0m40.648s<br>user 9m18.562s<br>sys 0m48.366s | real 0m54.010s<br>user 6m48.936s<br>sys 0m38.000s | real 0m32.341s<br>user 8m27.006s<br>sys 0m35.783s | real 5m46.422s<br>user 252m24.684s<br>sys 10m57.436s | real 0m49.982s<br>user 11m32.437s<br>sys 1m24.588s | real 1m11.182s<br>user 32m43.466s<br>sys 4m28.850s | real 1m3.163s<br>user 14m29.265s<br>sys 1m1.547s | real 1m0.202s<br>user 30m52.835s<br>sys 2m2.200s | real 1m11.161s<br>user 19m44.509s<br>sys 1m20.224s |
| T10 | real 0m35.810s<br>user 2m52.484s<br>sys 0m19.840s | **doesn't reconstruct** | real 2m1.935s<br>user 54m35.235s<br>sys 2m31.266s | **doesn't reconstruct** | real 0m38.576s<br>user 3m13.032s<br>sys 0m29.856s | **doesn't reconstruct** | real 2m29.201s<br>user 63m51.670s<br>sys 2m56.702s | **doesn't reconstruct** | real 0m15.244s<br>user 6m47.619s<br>sys 0m43.388s | real 0m19.543s<br>user 4m0.029s<br>sys 0m21.067s | real 0m58.627s<br>user 6m48.270s<br>sys 0m45.369s | real 0m26.452s<br>user 3m46.169s<br>sys 0m18.922s |
| T11 | real 0m57.154s<br>user 4m21.307s<br>sys 0m33.827s | real 0m28.594s<br>user 5m16.188s<br>sys 0m26.105s | real 4m0.837s<br>user 154m40.611s<br>sys 7m53.837s | real 0m34.613s<br>user 6m34.570s<br>sys 0m41.080s | real 0m57.218s<br>user 5m56.852s<br>sys 0m51.089s | real 0m28.512s<br>user 5m36.128s<br>sys 0m30.368s | real 4m42.793s<br>user 182m13.256s<br>sys 9m46.192s | real 0m46.138s<br>user 10m20.638s<br>sys 1m54.849s | real 0m50.356s<br>user 23m15.919s<br>sys 2m49.245s | real 0m56.233s<br>user 12m49.028s<br>sys 0m59.363s | real 1m53.462s<br>user 34m57.238s<br>sys 2m54.438s | real 1m13.915s<br>user 23m12.817s<br>sys 1m11.934s |
| T12 | elapsed 9:54.06<br>user 5666.09<br>system 201.66 | real 6m10.074s<br>user 106m34.685s<br>sys 4m17.712s | elapsed 53:23.97<br>user 146526.23<br>system 4842.22 | real 10m20.941s<br>user<br>170m13.594s<br>sys 10m45.900s | real 8m12.549s<br>user 97m25.751s<br>sys 3m53.114s | real 6m21.273s<br>user 108m26.167s<br>sys 4m30.485s | real 52m46.188s<br>user 2428m35.522s<br>sys 96m43.449s | real 9m21.252s<br>user<br>176m32.067s<br>sys 10m57.930s | real 13m32.707s<br>user 425m29.606s<br>sys 59m54.658s | real 11m54.151s<br>user 232m21.363s<br>sys 13m40.987s | real 14m49.488s<br>user 561m12.309s<br>sys 37m22.082s | real 14m21.677s<br>user 254m56.741s<br>sys 19m43.417s |
| T13 | real 8m15.889s<br>user 83m12.045s<br>sys 13m11.063s | real 8m42.431s<br>user 182m54.503s<br>sys 9m12.325s | elapsed 1:56:13<br>user 326985.97<br>system 8375.78 | elapsed 11:19.10<br>user 13948.33<br>system 1138.69 | real 8m1.042s<br>user 90m6.927s<br>sys 12m58.928s | real 7m53.973s<br>user 177m14.824s<br>sys 8m35.664s | real 118m1.693s<br>user 5646m6.506s<br>sys 129m34.099s | real 12m33.124s<br>user<br>344m44.170s<br>sys 20m13.202s | real 26m31.697s<br>user 915m27.772s<br>sys 38m48.730s | real 26m32.049s<br>user 872m25.418s<br>sys 38m59.208s | real 31m28.899s<br>user 1000m8.820s<br>sys 40m57.281s | real 28m39.551s<br>user 953m1.007s<br>sys 41m21.804s |
| T14 | real 22m20.461s<br>user 275m7.091s<br>sys 180m12.796s | elapsed 14:49.77<br>user 21174.71<br>system 1244.72 | elapsed 1:56:45<br>user 342283.42<br>system 9120.26 | real 16m30.784s<br>user<br>365m37.021s<br>sys 23m39.682s | real 22m22.005s<br>user<br>274m51.799s<br>sys 185m37.468s | real 14m48.525s<br>user 368m9.925s<br>sys 28m17.028s | real 117m45.300s<br>user 5734m4.810s<br>sys 153m24.307s | real 17m49.536s<br>user<br>418m57.785s<br>sys 27m35.760s | real 15m26.790s<br>user 542m56.410s<br>sys 38m58.651s | real 36m29.920s<br>user 388m37.709s<br>sys 27m47.447s | real 24m13.578s<br>user 986m33.831s<br>sys 59m42.900s | real 29m26.902s<br>user 615m48.642s<br>sys 40m45.544s |

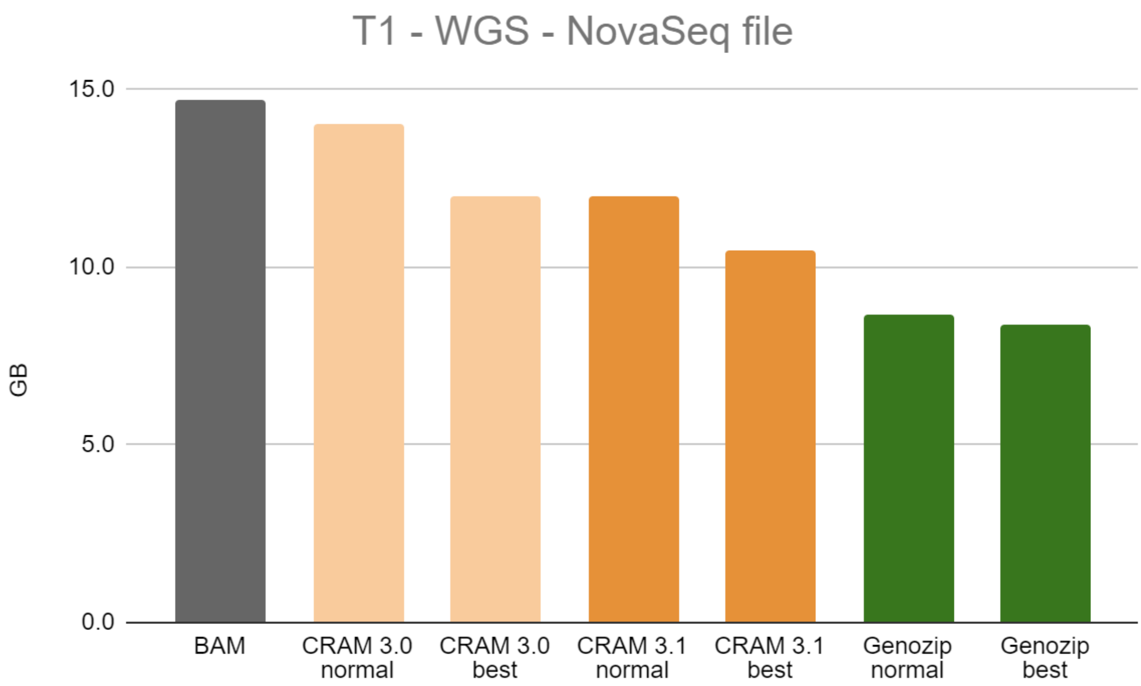


Figure S1: Benchmark file T1

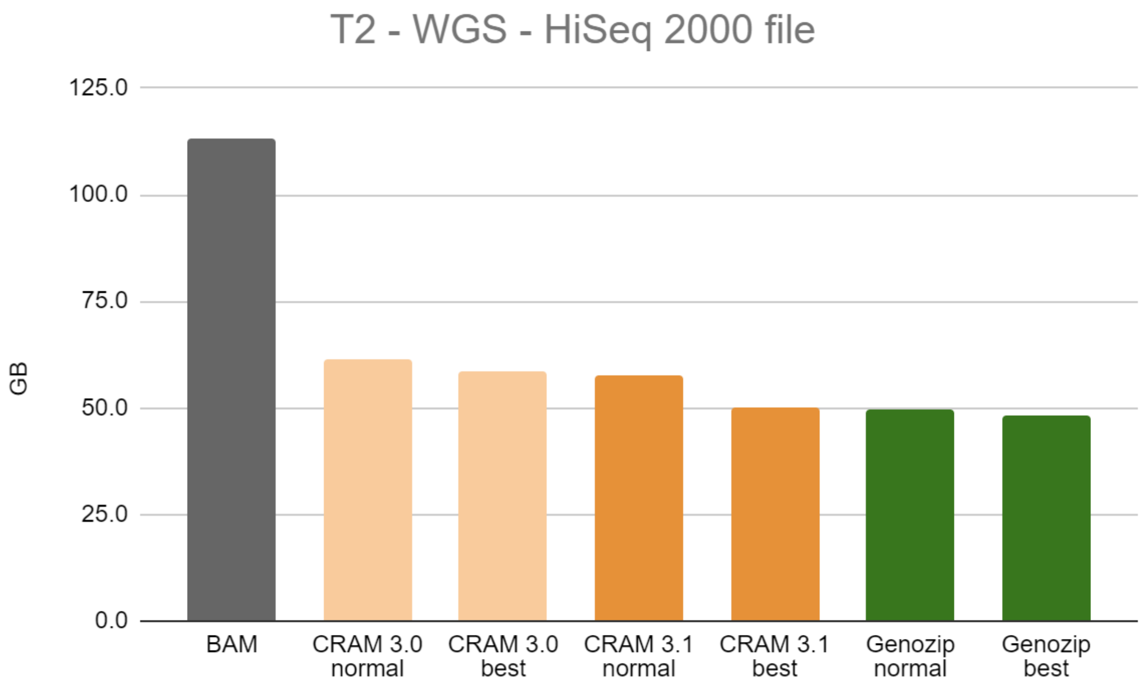


Figure S2: Benchmark file T2

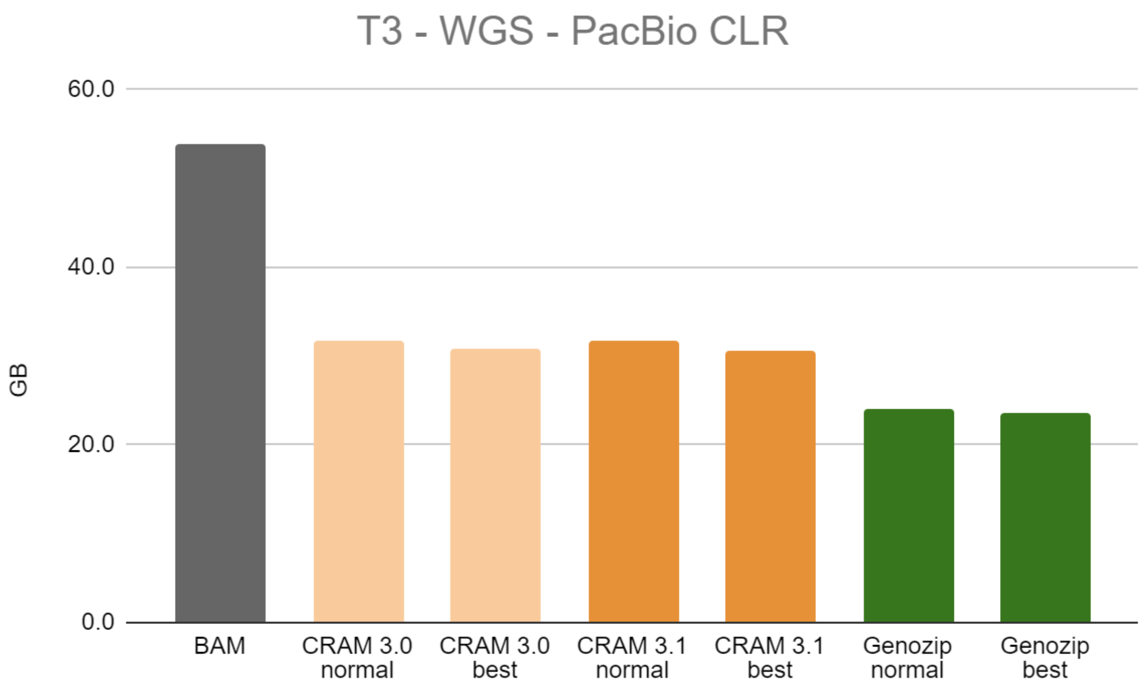


Figure S3: Benchmark file T3

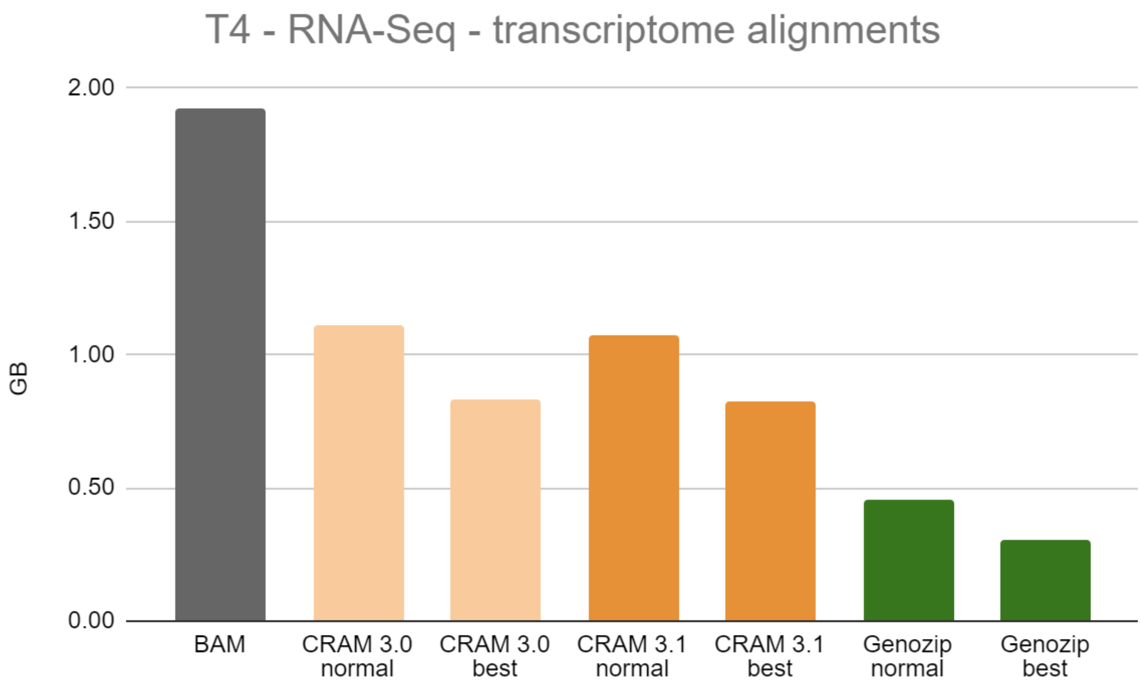


Figure S4: Benchmark file T4

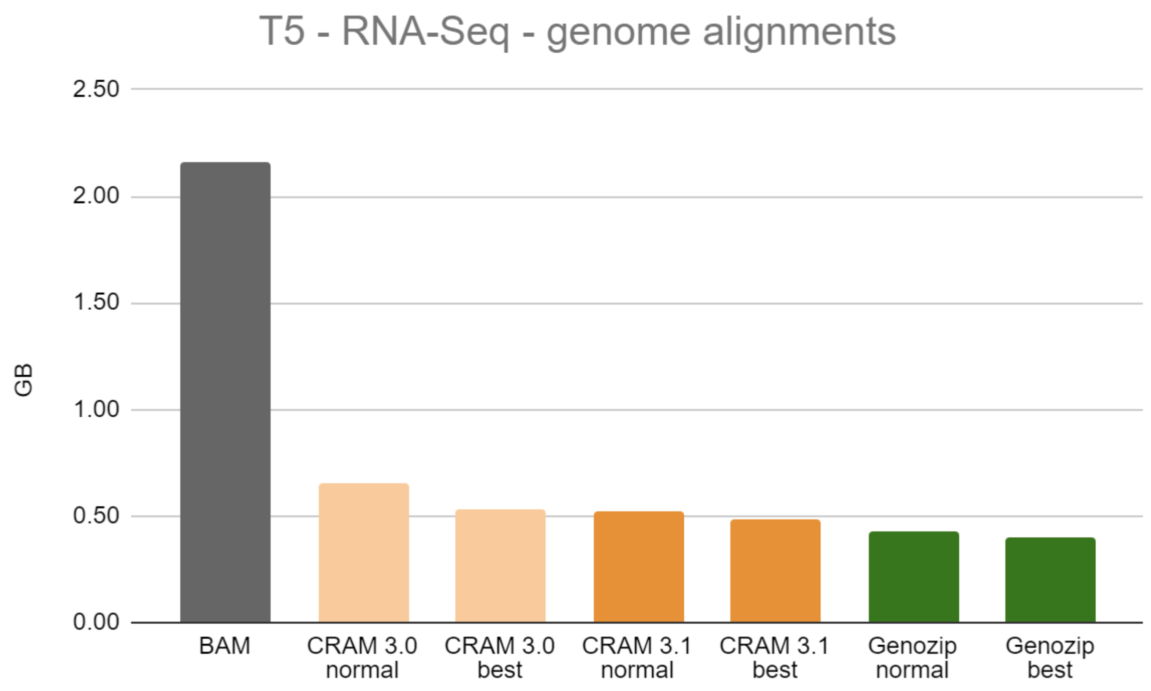


Figure S5: Benchmark file T5

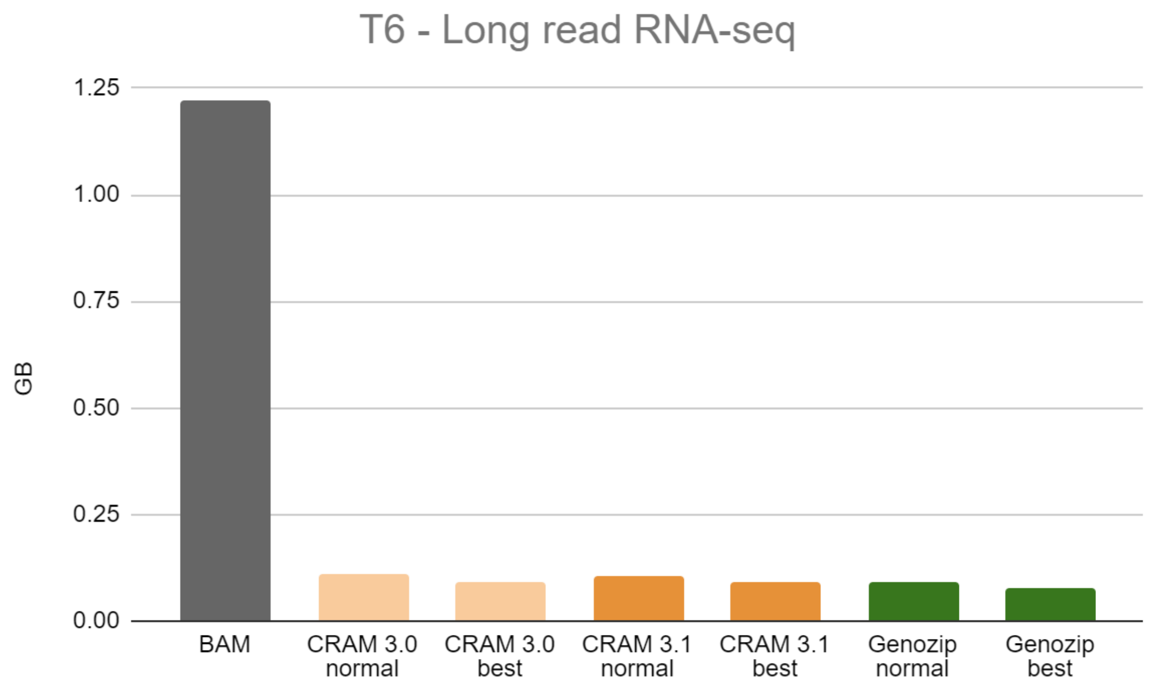


Figure S6: Benchmark file T6

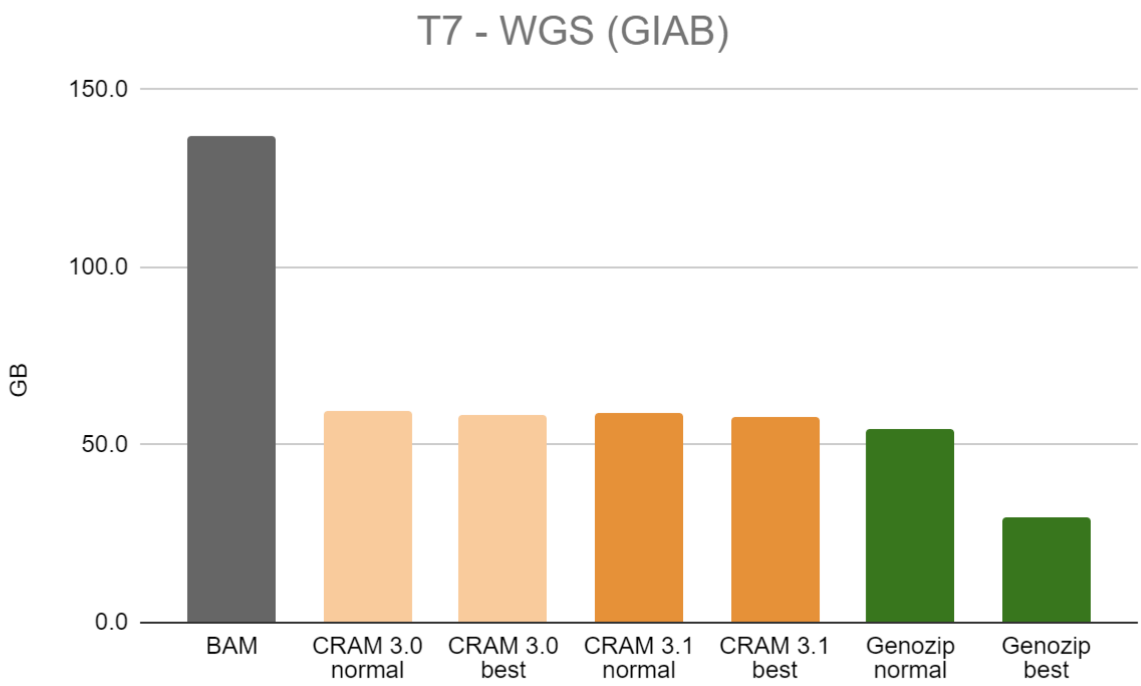


Figure S7: Benchmark file T7

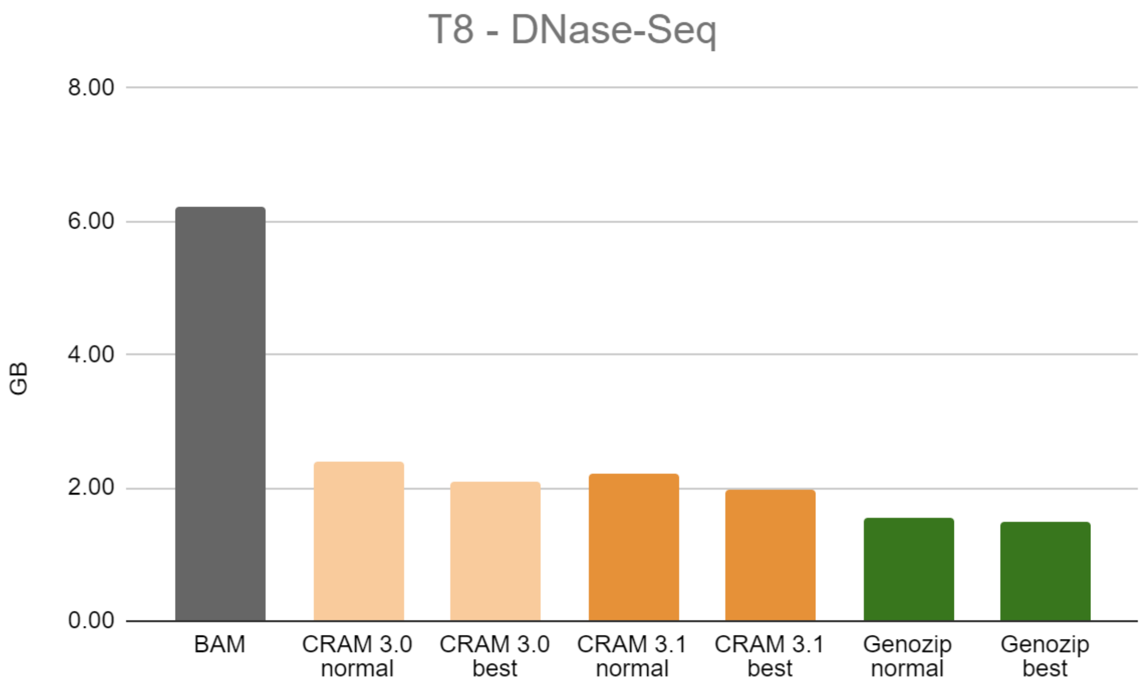


Figure S8: Benchmark file T8

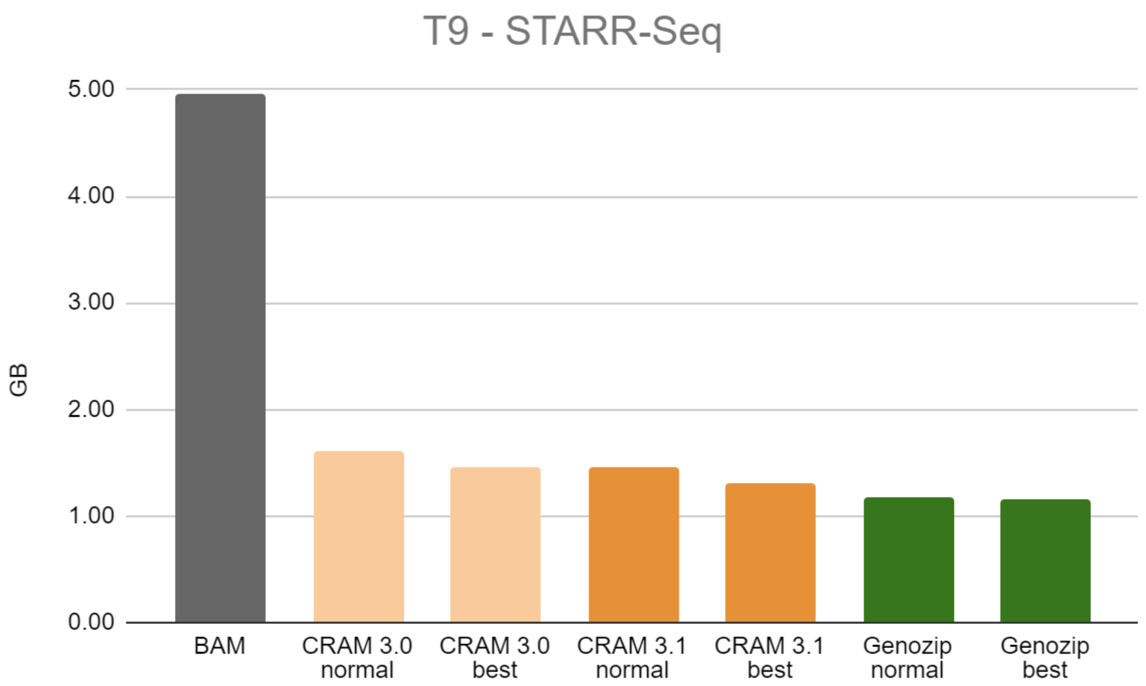


Figure S9: Benchmark file T9

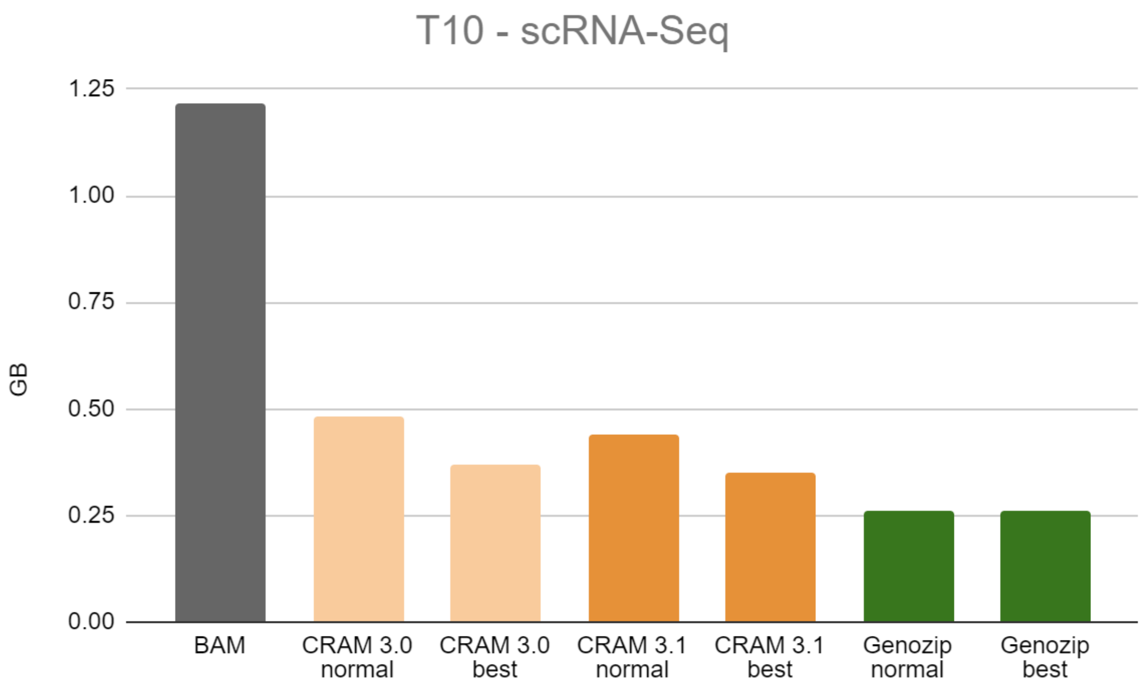


Figure S10: Benchmark file T10

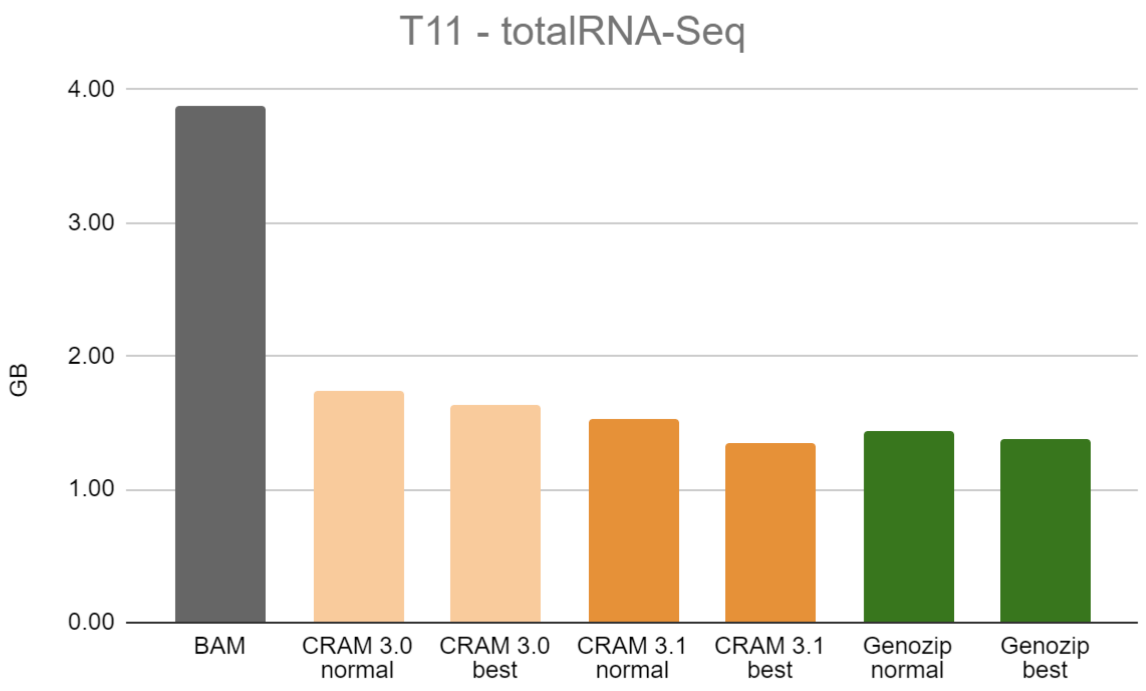


Figure S11: Benchmark file T11

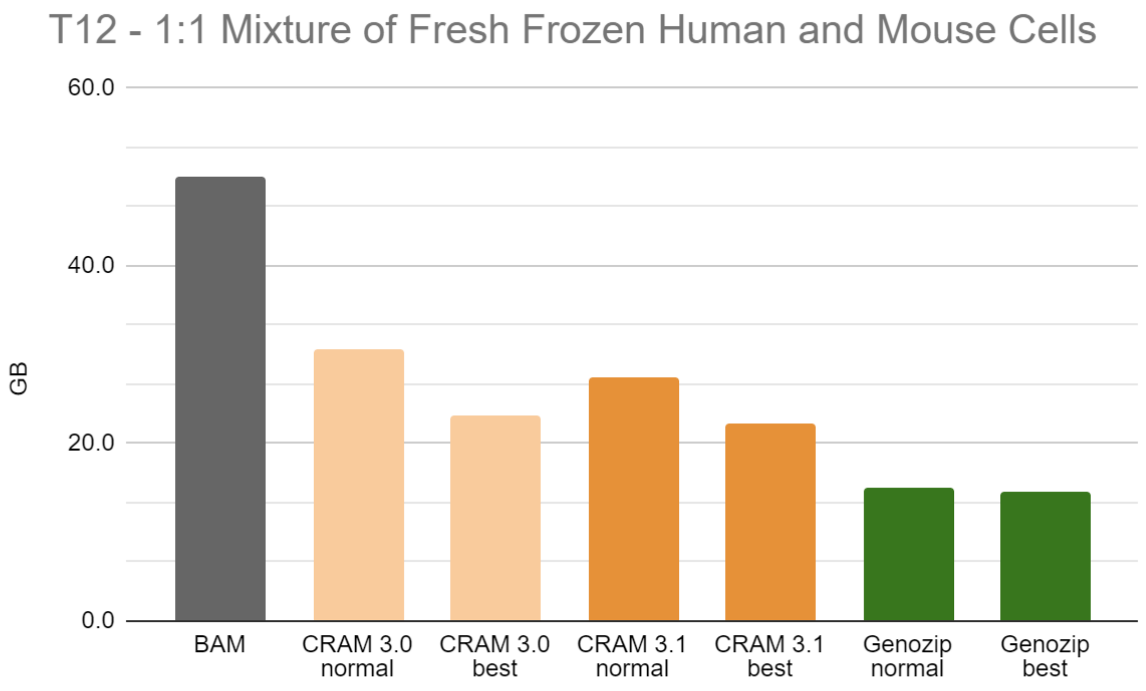


Figure S12: Benchmark file T12

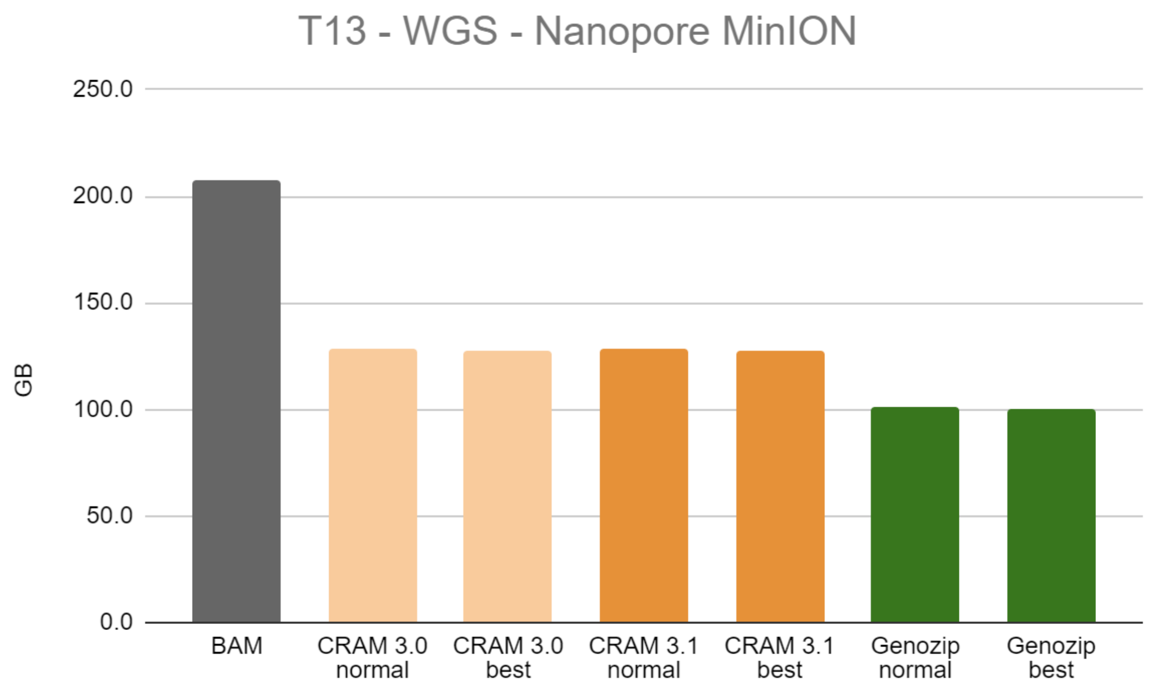


Figure S13: Benchmark file T13

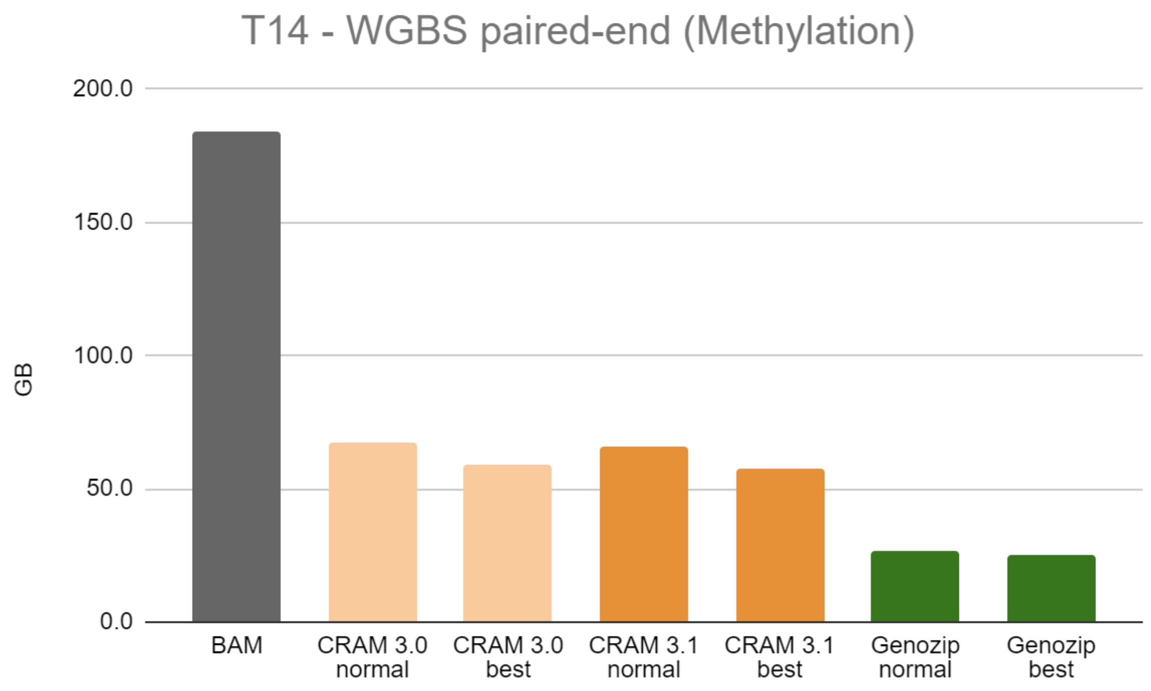


Figure S14: Benchmark file T14
